# Cross-feeding enables robust coexistence between four bacterial species

**DOI:** 10.64898/2026.04.20.719729

**Authors:** Snorre Sulheim, Miguel Teixeira, Eric Ulrich, Alisson Gillon, Samuele E. A. Testa, Prajwal Padmanabha, Daniel Machado, Sara Mitri

## Abstract

Microbial diversity is often assumed to be limited by the number of available resources, yet many communities persist well beyond that expectation. Understanding the mechanisms that enable such coexistence remains a central question in microbial ecology. Here, using a four-species bacterial consortium, we asked whether coexistence can emerge from interactions between species rather than from the external environment alone. Across 31 simple nutrient conditions, including 16 single-resource environments, all four species persisted and repeatedly reached stable coexistence. We then chose 27 additional conditions to further probe the boundaries of coexistence by varying resource concentrations, temporal dynamics, nutrient complexity and relief of auxotrophy-associated dependencies, and only observed the extinction of one species in one of these conditions.

Although the community composition in each environment was largely shaped by species’ fitness on the supplied resources, experimental assays and consumer-resource modeling showed that the coexistence was not explained by resource supply, but rather by cross-feeding and niche partitioning of metabolic byproducts. These metabolic interactions were strong enough to sustain coexistence even for species unable to use the supplied resources directly. Furthermore, robust coexistence across environments appears to be an emergent property of microbial communities, ingrained in members’ metabolic byproduct profiles and niche differences. Our findings demonstrate how microbes can increase the chemical complexity of their environment sufficiently to maintain coexistence well beyond what is expected from external resource supply.

**Significance:** Understanding the drivers of microbial diversity is essential for managing natural ecosystems and designing synthetic microbiomes. This study challenges the conventional application of the competitive exclusion principle, demonstrating that a four-species consortium can coexist across 31 chemically and metabolically diverse one- and two-carbon source environments. By systematically testing and ruling out alternative stabilizing mechanisms, we show that co-existence is an emergent property of the consortium, sustained by metabolic cross-feeding and niche partitioning. Guided by computational models, we identify hallmarks of robust co-existence in simple environments, including high variance in resource affinities and growth on partner-derived metabolites. Our work demonstrates how microbes modify their environment to sustain high diversity and provides principles for designing synthetic microbiomes that persist across environments.

## Introduction

The stunning microbial diversity found in natural environments has impressed microbiologists since the 17th century. ^1–5^ This diversity – as with non-microbial ecosystems^6,7^ – correlates positively with the stability and functions of microbial ecosystems. ^8^ In the human gut, for instance, diminished microbiome diversity is associated with recurrent *Clostridium difficile* infections,^9^ while in terrestrial systems, soil microbiome diversity drives plant productivity and nutrient retention. ^10^ Understanding when and why microbes coexist is therefore critical for managing ecosystem stability and function.

Theoretical expectations of diversity are governed by the competitive exclusion principle, which posits that the number of coexisting species in a well-mixed environment cannot exceed the number of limiting resources. ^11,12^ However, empirical observations from nature and laboratory experiments frequently reveal richness that exceeds these theoretical bounds.^13–18^ Several mechanisms have been proposed to explain this excess diversity, including environmental fluctuations combined with variation in growth traits, favoring different species at different times,^15,19^ predation or inhibition of the dominant species,^20,21^ cross-detoxification^22,23^ or cross-feeding of metabolites. ^24–27^ Central to most of these mechanisms is negative frequency dependence, whereby rare genotypes have a growth advantage over the more common ones, leading to stable equilibrium frequencies.

Despite these insights, it remains challenging to predict when species will coexist or not, even in simple environments, ^28–30^ and it is unclear whether coexistence is largely determined by the number and identity of the supplied resources or whether it is a result of species interactions. Bridging this gap requires assembling a single well-characterized community in diverse environments and quantifying the abundances of its members across several orders of magnitude to discern between extinction and persistence of low-abundance species.

In this study, we conduct such experiments using a synthetic four-species community and demonstrate that these taxonomically diverse bacteria coexist consistently even when provided with only a single carbon source. We rule out alternative stabilizing factors as explanations for the observed coexistence, including auxotrophies and resource fluctuations. Rather, our findings reveal that microbes actively increase the chemical complexity of their environment through niche construction, maintaining stable coexistence via cross-feeding beyond the levels predicted by external resource supply. In other words, the competitive exclusion principle remains valid: What was seen as a paradox – that more species can coexist than externally supplied resources – arises because the supplied limiting resources are far fewer than the total number of limiting resources available following microbial growth.

## Results

### A synthetic four-species community robustly coexists across simple nutrient environments

To test the limits of the competitive exclusion principle (CEP), we utilized a synthetic model community comprising four taxonomically and functionally distinct bacteria: *Agrobacterium tumefaciens* (At), *Comamonas testosteroni* (Ct), *Microbacterium liquefaciens* (Ml), and *Ochrobactrum anthropi* (Oa). These strains were originally isolated from industrial machine oils, ^31^ and have since been used as a model system to study synthetic communities. ^22,23,32^ A critical advantage of this consortium is the ability to resolve individual species abundances on selective agar media even at low frequencies,^23^ essential for distinguishing between competitive exclusion and species declining to a low-abundance steady-state. These species, isolated in the past from toxic metal-working fluids based on their ability to survive in and degrade such environments, ^22,33^ possess varying metabolic capabilities and nutritional dependencies.^34,35^ Notably, At and Ct are prototrophs while previous experiments have shown that Oa and Ml depend on co-cultivated partners for prolonged growth. ^23^ To characterize these dependencies, we first identified putative auxotrophies based on the absence of essential biosynthetic enzymes in the annotated proteomes, and then conducted drop-out or drop-in assays to confirm or reject these metabolic dependencies. Gaps in vitamin biosynthesis pathways matched *in vitro* experiments, verifying that Oa is a thiamine auxotroph and Ml is auxotrophic for both biotin and thiamine (Fig. S1A-C). In contrast, predicted amino acid auxotrophies of Ml did not match *in vitro* results, which revealed an absolute dependency on exogenous cysteine. Furthermore, while genetically prototrophic for proline, supplementation was required to bypass a substantial 48-hour lag phase (Fig. S1D-E). By incorporating these natural auxotrophies, this four-species community serves as a tractable model community that captures the functional dependencies found in natural microbiomes.^36^

We characterized the growth of each species across 33 metabolically and chemically diverse carbon sources (Fig. S2A-B), revealing distinct niche differences and a trade-off between versatility and growth rate (Fig. S2C-D). Ct was the least versatile, unable to utilize 14/33 substrates (including all sugars), but frequently exhibited the fastest growth on organic acids and amino acids. In contrast, Ml utilized 32 of 33 tested carbon sources but grew slowly in most conditions. From this set, we selected 16 carbon sources to construct 32 resource environments for our initial community assembly experiment, consisting of 16 single-resource and 15 paired-resource, and a no-carbon control condition. These substrates included organic acids, sugars, amino acids, an alcohol (glycerol), a sugar-alcohol (myo-inositol) and a nucleoside (inosine), and all environments were normalized to an equal amount of carbon (90 mM C). The community was then subjected to eight growth-and-dilution cycles, each lasting 72h in these different media. By selecting a chemically and metabolically diverse array of carbon sources, as well as choosing carbon sources favoring different species, we sought to maximize environmental variability and test the boundaries of coexistence.

Given the limited number of provided carbon sources, our expectation was that competitive exclusion and low richness would be prevalent. Instead, all four species coexisted in every environment tested, reaching stable steady-states by the eighth transfer and often already after four or five transfers (Fig. 1A and B). Even species unable to utilize the provided carbon sources were able to reach steady-state population sizes, in some environments up to 10^8^ CFUs/mL, suggesting that whatever allows these species to grow is sufficient to sustain such population sizes and an average growth rate above the ∼0.064h^-1^ effective dilution rate imposed by the 1/100 dilution every 72h (citrate and inosine environments diluted 1/10, see Methods). All species except Ml were also still present at the end of the experiment in the no-carbon control, although at significantly lower abundances than in the other environments (*P* < 1e-3 for all species and conditions, see Methods). Low-level growth in no-carbon conditions is a well-known challenge in microbial cultures that has been shown to support low-abundance communities in similar experiments previously. ^18^

**Figure 1:**
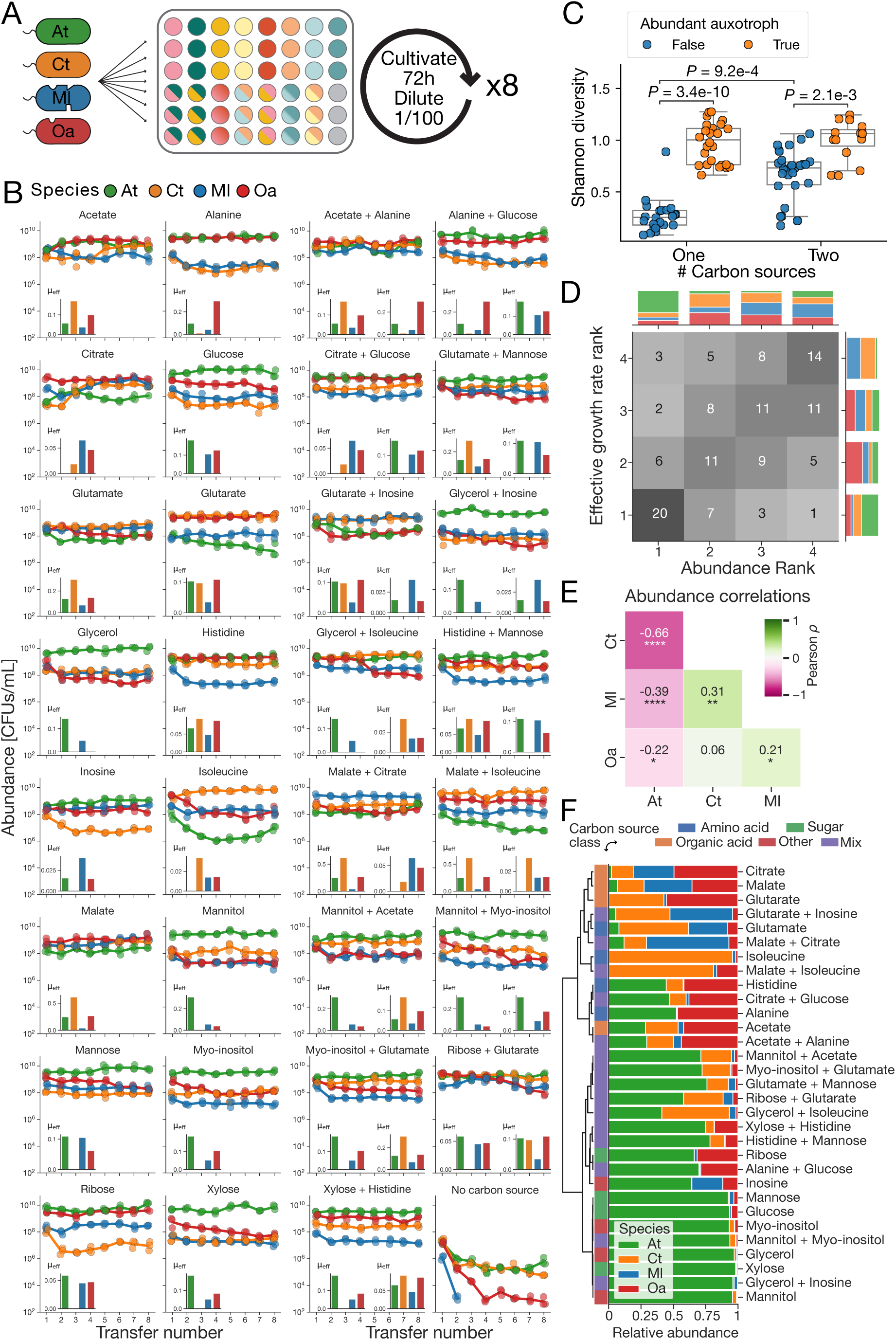
Four species coexist across 31 conditions. A) Schematic illustration of the community assembly experiments. B) Quantified species abundance at the end of each of the 8 consecutive batch transfers. Single carbon source environments are shown in the two leftmost columns, while environments with two carbon sources and the no-carbon control are shown in the two rightmost columns. The inset bar charts show the effective growth rates for each species measured from separate monoculture experiments, and for the two-carbon source environments left and right inset corresponds to first and second carbon source, respectively. C) Environments with an abundant auxotroph (Oa or Ml is >25% of the total population) have increased diversity (Fig. S3B), but this increase is only significant for the single carbon source environments. For the environments without an abundant auxotroph, the diversity is significantly higher when two carbon sources are provided. Statistical tests: Kruskal-Wallis followed by Dunn’s posthoc test with Benjamini-Hochberg correction. D) Community assembly abundance ranks are highly predictable from monoculture effective growth rates, with 84.7% (105/124) of observations falling within ±1 of the predicted rank. E) Pearson correlations of species’ abundance in the community assembly experiment (\**P* < 0.05, \*\**P* < 0.01, \*\*\**P* < 0.001, \*\*\*\**P* < 1e-4). F) Final relative abundance in the community assembly experiments (mean of three replicates over the last two time points clustered by similarity in community composition).

Contrary to the expectations of the competitive exclusion principle, Shannon diversity was not primarily driven by the complexity of the supplied resources (*P* = 0.21, Mann-Whitney U; *n*_1_ = 48, *n*_2_ = 45, Fig. S3A). Instead, the prevalence of auxotrophic members emerged as the principal determinant of community structure. Environments where an auxotroph (Oa and Ml) constituted more than 25% of the total population showed a significantly higher diversity (*P* = 7.1e-11, Mann-Whitney U, Fig. S3B). By partitioning the data by both resource complexity and auxotroph prevalence, we observed that auxotroph prevalence had the strongest influence on single-carbon environments (Fig. 1C). In the absence of an abundant auxotroph, increased resource complexity did increase Shannon diversity (Fig. 1C), but this effect was secondary to the stabilizing influence of metabolic dependency. This phenomenon is likely driven by the inherent constraints of cross-feeding: the reliance of auxotrophs on partners for essential vitamins or amino acids effectively caps their relative abundance, often resulting in two or more species persisting at nearly equal frequencies even on a single substrate (see e.g., alanine or glutarate environments, Fig. 1B).

We next investigated the extent to which species’ rank abundances in the assembled communities could be predicted from their growth in monoculture. Monoculture growth rate ranks were significantly correlated with final abundance ranks (Kendall’s *τ* = 0.33, *P* = 1.3e-5, N = 124), but predictive accuracy improved when using “effective” growth rates that also accounted for lag and yield differences (Kendall’s *τ* = 0.46, *P* = 1.2e-9, N = 124, Fig. 1D, see Methods). The most abundant species was predicted with higher fidelity than subordinate members, likely reflecting that while primary resource utilization governs dominance, the relative abundances of subordinate species are also shaped by secondary processes, such as metabolite cross-feeding, that cannot be informed by monoculture growth data. To interrogate these putative interactions, we analyzed species abundance correlations, recognizing that such associations integrate the net effects of both nutrient competition and cross-feeding. While At was negatively correlated with all other community members, Ml’s abundance was positively correlated with Oa and Ct (Fig. 1E). The negative correlations involving At reflect its competitive dominance in the environments where it exhibits the highest effective growth rate (sugars), where it frequently exceeds more than 90% of the population suggesting that it provides little support for co-cultured partners (Fig. 1F). In contrast, the positive correlations in abundances of Ml with Ct and Oa were not mirrored by correlations in effective growth rate (Fig. S3D). This decoupling suggests their positive association is not driven by shared resource preferences, but is rather an emergent result of mutualistic or commensal cross-feeding interactions.

### Coexistence is robust to environmental perturbations

To investigate the mechanisms underlying the unexpected and consistent excessive richness, we systematically tested several ecological hypotheses that might explain coexistence beyond the competitive exclusion principle. Given that auxotrophic dependencies appeared to shape community structure, we first hypothesized that these requirements were the primary stabilizing force. We therefore repeated the community assembly experiment with seven of the same single carbon source environments, but now also supplemented with the amino acids (cysteine and proline) and vitamins (thiamine and biotin) needed by Oa and Ml. Relieving these metabolic dependencies were not sufficient to break coexistence nor change the average diversity (Fig. 2A-B, Fig. S4A), but we found a reduced, and no longer significant, increase in Shannon diversity in the environments with an abundant auxotroph (>25%, *P* = 0.11, Mann-Whitney U, Fig. S4B). Note that both proline and cysteine can also be used as carbon sources, and all four species also coexisted when only provided the amino acid and vitamins supplements (Fig. S4C). As expected by their metabolic requirements, Oa reached higher steady-state densities in supplemented conditions, and Ml showed a similar expansion in inosine, the only carbon source on which it exhibits the highest effective growth rate. Despite these shifts in individual species performance, the persistence of all four species across all supplemented environments suggests that the underlying stabilizing mechanisms extend beyond simple vitamin or amino acid dependencies.

**Figure 2:**
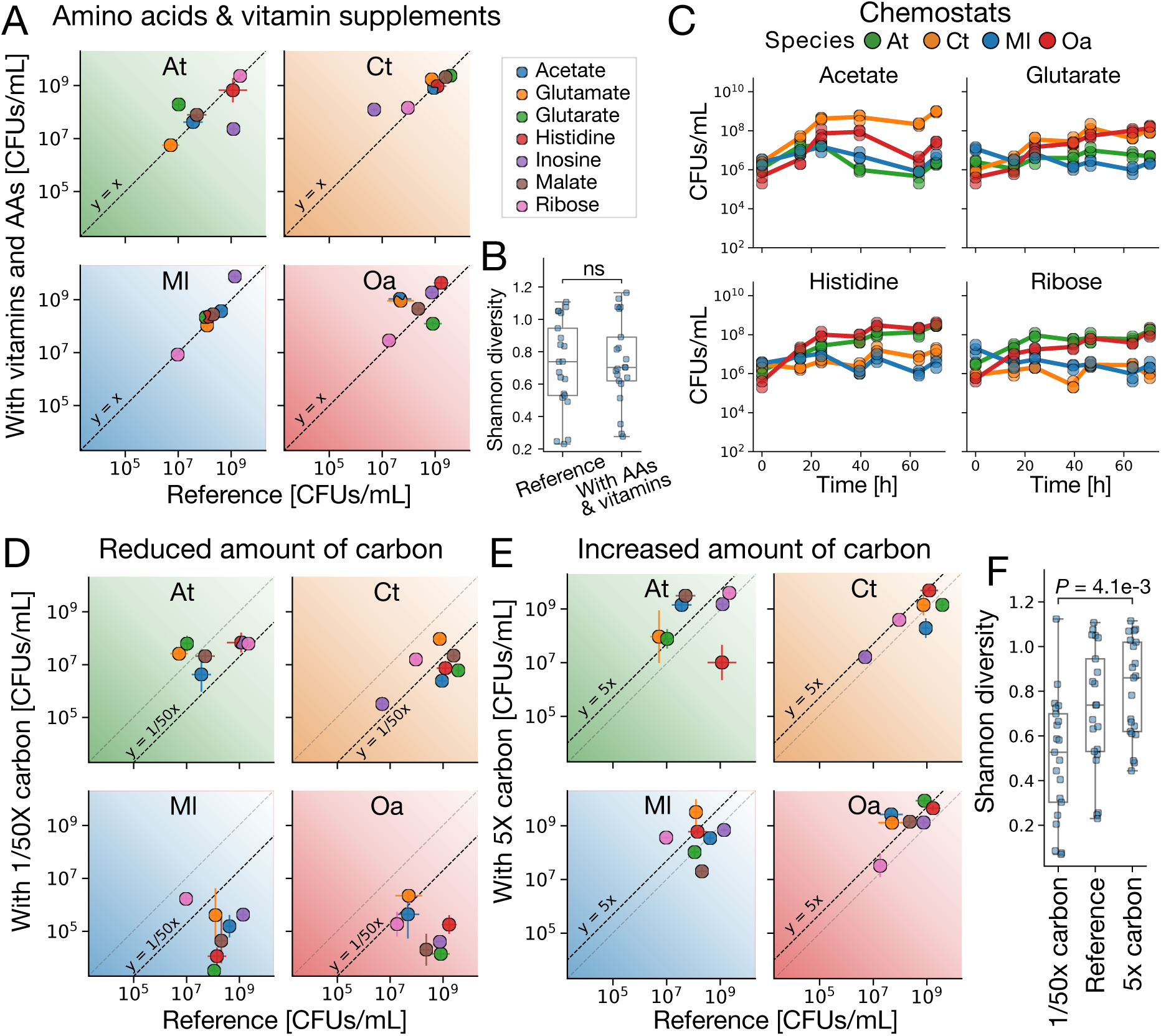
Coexistence is robust to different environments. A) Effect of adding vitamins (thiamine and biotin) and amino acids (proline and cysteine) to seven reference single carbon source conditions, here shown as final population sizes with (y-axis) or without (x-axis) supplements after eight transfers (mean ± standard deviation, *n*=3). B) We find no significant difference in Shannon diversity between the reference conditions and the environments with additional vitamins and amino acids (*P* = 0.72, Mann-Whitney U). C) Continuous cultivation in chemostats did not disrupt species coexistence, despite considerably lower abundances (Fig. S4C). D) All four species also coexisted with a 50-fold reduction in carbon, but these conditions seemed to disfavor the auxotrophs (Ml and Oa) with abundances in most cases well below the naive expectation of 50X less than in the reference conditions. E) Ct went immediately extinct in the 5X malate condition. Otherwise all four species coexisted, but with significant shifts in abundances compared to the reference condition. F) We found a significant difference in Shannon diversity between the 1/50X and 5X carbon conditions (Kruskal-Wallis H-test followed by Dunn’s posthoc test, Benjamini-Hochberg correction)

We next addressed whether the periodic resource fluctuation in the consecutive batch cultures could explain the observed coexistence. Theoretical studies have shown that resulting temporal fitness advantages can support increased species coexistence, ^37–40^ for example due to trade-offs between growth rate and lag-time^41^ or growth rate and resource affinity. ^42^ To test this, we repeated the community assembly experiment in four single carbon source environments in chemostats with continuous inflow of nutrients and outflow the microbial culture. Species abundances increased in the chemostats to reach steady-states within 39 hours, with all species coexisting in all four conditions (Fig. 2C). The steady-state abundances in chemostats were correlated with the steady-state abundances in the batch transfer community assembly (Pearson *ρ* = 0.67, *P* = 4e-3), but abundance ranks shifted compared to the batch cultures and abundances were consistently lower in the chemostat experiment (Fig. S4C), reflecting the transition from favoring species with high maximum growth rate and short lag-time in batch cultures to favoring the species able to sustain the imposed dilution rate at the lowest resource concentration in the chemostats.^14^

Finally, we explored how interaction strength influenced coexistence. Previous research has demonstrated the importance of interaction strength for the stability and diversity of microbial communities, and that interaction strengths can be modulated by total resource availability. ^43,44^ We therefore repeated the community assembly experiment with 5-fold increase (5X) or a 50-fold reduction (1/50X) in carbon. Except in the 5X malate condition, where Ct disappeared after the first transfer likely due to toxic substrate concentrations, all species coexisted in both the high and low resource concentrations (Fig. 2D and E, Fig. S4A). Community compositions were, however, strongly affected by the change in resource concentration, potentially reflecting rate *vs* resource affinity differences, toxic effects in the 5X conditions,^44^ or differences in metabolic byproducts and cross-feeding patterns. We observed significantly lower diversity in the 1/50X environments compared to the 5X environments (Fig. 2F), contrasting with previous results showing increased diversity in low nutrient environments. ^43,44^ Interestingly, the low concentrations seemed to disfavor the auxotrophs, with most abundances below the naive expectation of 1/50 of the abundance in the reference condition, potentially suggesting that cross-feeding is weaker when resources are scarce. While a general correlation between growth rate and metabolite release remains undescribed, overflow metabolism causing the release of fermentation end-products such as acetate indeed only occurs when resources are abundant and the growth rate is high. ^45^ Finally, we also found that the four species coexisted in rich media (LB and TSB), with high diversity in TSB and very low diversity in LB, with Ml almost going extinct and reaching abundances comparable to our negative control without any provided carbon source (Fig. S4D). Collectively, these results demonstrate that the observed four-species coexistence is remarkably robust and resilient to changes in nutrient complexity, temporal dynamics, and resource concentration.

### Cross-feeding explains coexistence

Having excluded alternative explanations for the observed coexistence, cross-feeding emerged as the primary stabilizing mechanism. Microbial metabolite release is common and chemically diverse in *in vitro* cultures, ^46–48^ and has been linked to coexistence both on the strain and species level in similar experiments. ^16,18,49^ To calibrate our expectations for how cross-feeding shapes coexistence, we employed a consumer-resource modeling (CRM) framework, which explicitly describes the coupled dynamics of species and resources. ^50,51^ Building on previous work demonstrating that cross-feeding can promote coexistence and that CRMs capture relevant patterns of microbial communities, ^18,52^ we used this framework to quantify the influence of specific microbial traits on community richness. Our CRM simulates growth, resource depletion and accumulation as a function of species’ resource affinities, maximum uptake rates and metabolite release profiles and thus captures both resource competition and cross-feeding (Fig. 3). We began by screening random four-species communities on a single provided resource with an increasing number of metabolic byproducts and increasing variability in resource affinities at three levels of metabolic versatility, defined as the average number of utilizable resources for each species. High or low metabolic versatility can be intuitively understood as communities composed of mostly generalist or specialist species, respectively. Our simulations revealed that community richness depends strongly on the interplay between resource versatility and variation in resource affinities (Fig. 3B). Intuitively, reduced versatility combined with high affinity variation increases the probability of each species specializing on a “private” metabolic niche, thereby promoting stable coexistence. Even at moderate number of byproducts and moderate resource affinity variation (below the standard-deviation of measured, log-transformed resource affinities of 1.6^53^), the model frequently produced stable four-species assemblages on a single primary resource. When the number of byproducts and variation in resource affinities were large, and metabolic versatility low, stable 4-species communities were common (Fig. 3B).

**Figure 3:**
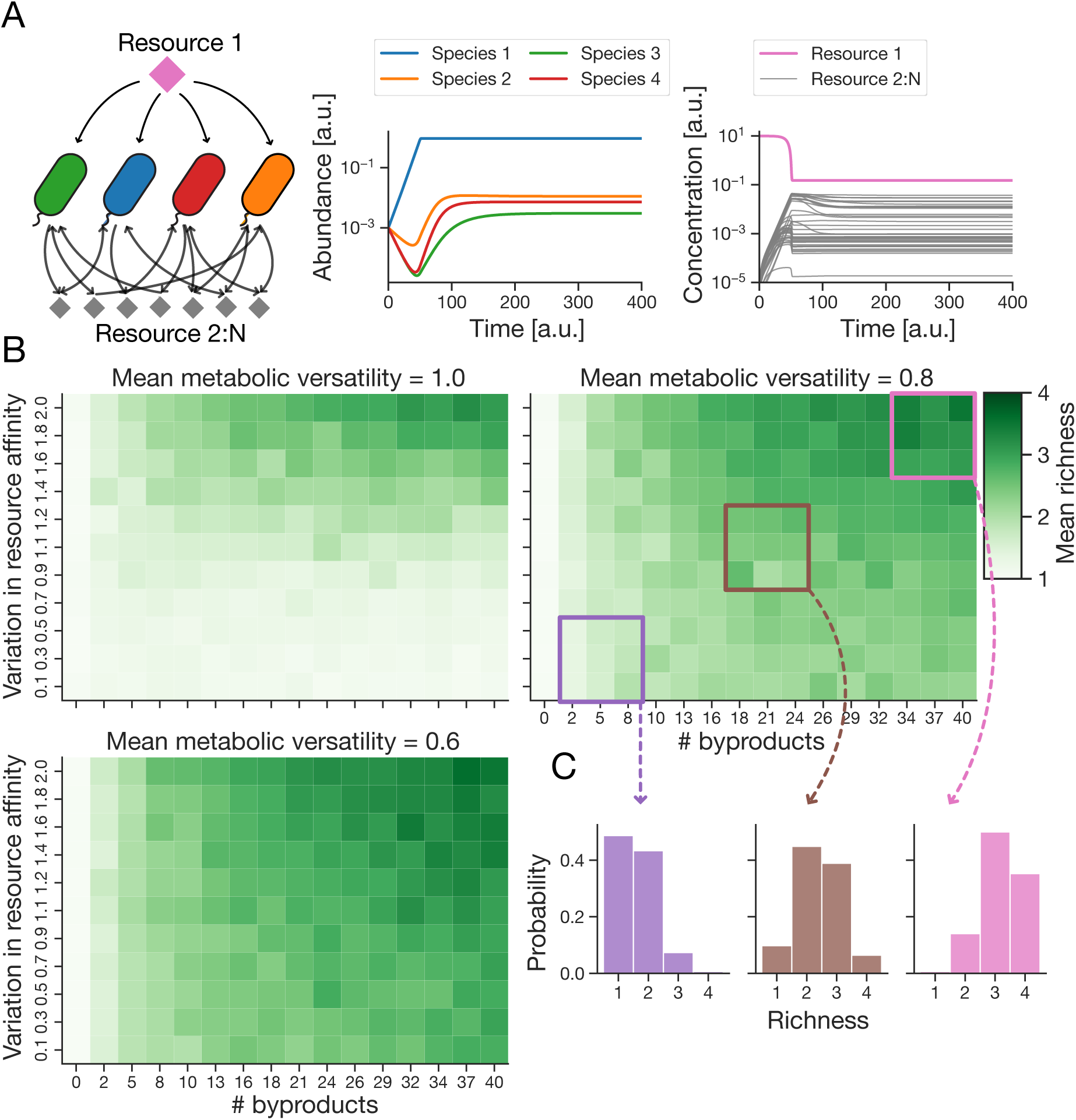
A consumer-resource model explains coexistence on a single resource. A) Illustration of the consumer-resource model and an example simulation showing the simulated species and resource dynamics leading to coexistence. B) The heatmaps display mean richness across 50 simulations for each combination of parameters. The variation in resource affinity is the standard deviation of the log-normal distribution used to draw half-saturation constants. The mean metabolic versatility is the probability of each species being able to utilize each resource. C) The distribution of richness across parameter regions indicated in panel B.

To better understand the properties determining a community’s ability to maintain high richness across environments, we conducted a new set of simulations across a parameter subset expected to yield a diverse range of richness outcomes. Rather than drawing new species pools (with new species properties) for each simulation, we iteratively simulated growth across seven different resources for each community (by changing the index of the single, provided resource in our CRM), and asked whether the richness of some communities was robust to the identity of the provided resource. Indeed, we found a significant correlation between the richness of a community grown on two different resources (Fig. 4A). To extend beyond pairwise comparisons, we quantified the conditional probability of a community maintaining its richness across *n* sequential resources. Consistent with the pairwise results, the conditional probability increased substantially with the number of preceding resources yielding the same richness (Fig. 4B). Essentially, communities that exhibited high richness in one environment tended to exhibit high richness across other environments, demonstrating that richness is largely an intrinsic property of the community itself, emerging from the network of metabolic interactions rather than the identity of the external resource supply. In contrast, species’ abundances, or abundance ranks, were not correlated between two supplied resources (Fig. S5A-B). Rather, species’ abundance ranks were largely determined by their fitness on the provided resource (Fig. S5C), which was a good predictor of the most abundant species, largely reflecting our experimental findings (Fig. 1D).

**Figure 4:**
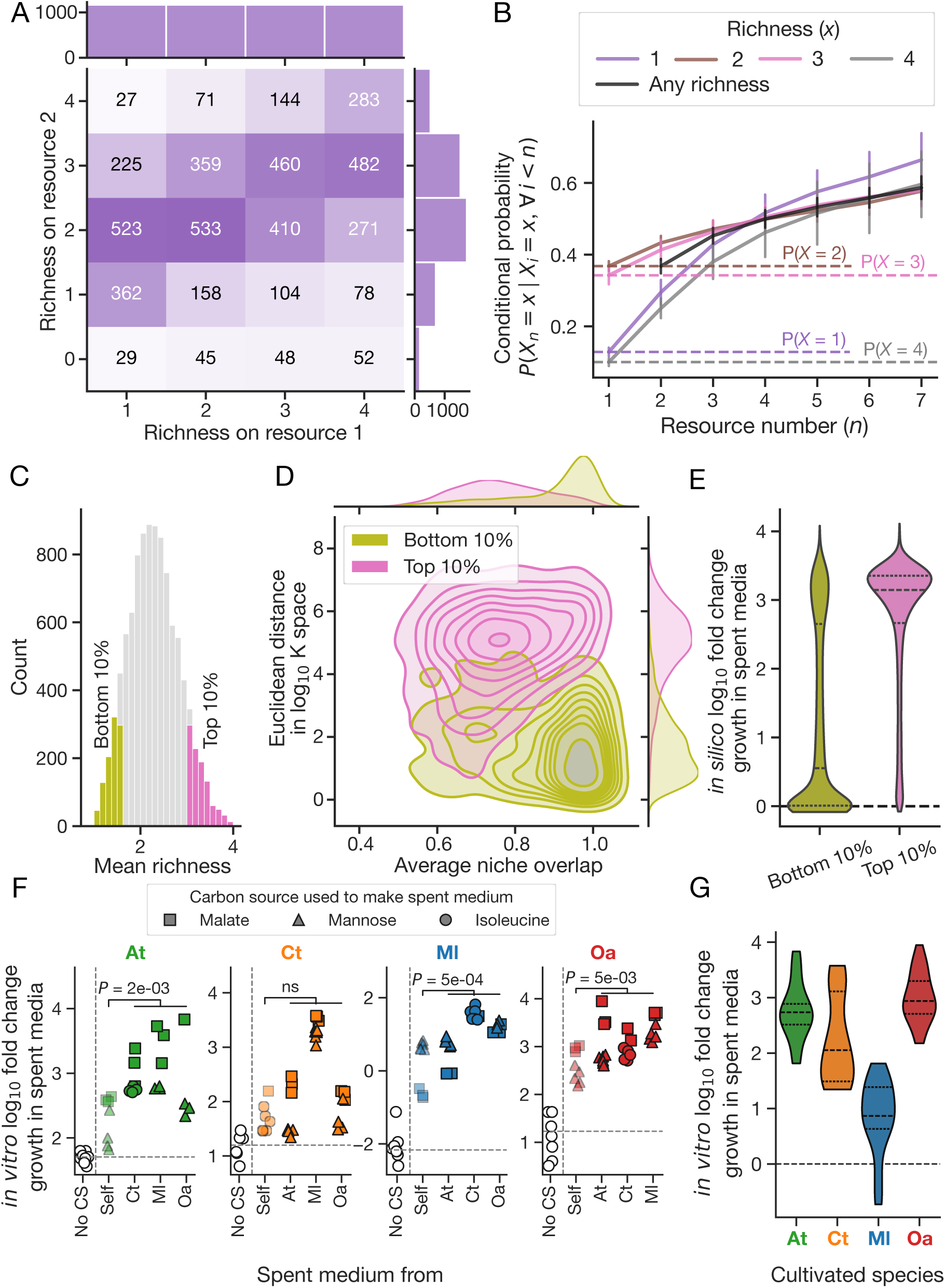
Consumer-resource models reveal hallmarks of robust community richness. A) Richness of CRM communities is highly correlated between provided resources, as displayed here for resource 1 and resource 2. This data was sub-sampled to a uniform distribution of richness outcomes from 1 to 4 on resource 1 to simplify interpretations. B) Conditional probability of maintaining a specific richness on resource, given consistent richness across all preceding resources. The black line represents the chance of having the same richness on resource *n* as on all preceding resources. The dashed lines represent the unconditional probability of each richness. C) Distribution of mean richness across seven provided resources for CRM communities, highlighting the most rich (top 10%) and least rich (bottom 10%) used to further understand the hallmarks of consistently rich communities. D) The least and most rich CRM communities are well separated on the euclidean distance in log_10_ K space and average niche overlap, K denoting the vector of resource affinities (Methods). E) In contrast to the least rich communities, the top 10% most rich communities consistently achieve high growth yields when cultivated on simulated spent media. F) *In vitro*, we also found that the species in general grew well in spent medium from one of the other species, and even in its own spent medium, here shown as log_10_ fold change in cell population [CFUs/mL] from inoculation to end of batch culture (after 72h). Statistical tests of growth in own spent media vs spent media from partners using Mann-Whitney U. G) Violin plots summarizing the distribution of *in vitro* growth in spent media for each of the four species (summarizing data in panel F), qualitatively recapitulating CRM simulations in spent media for the most rich communities shown in panel E.

To identify the metabolic properties of consistently rich communities, we calculated the average richness across the seven resources and compared the top 10% with the bottom 10% (Fig. 4C). In the parameter space, these two groups were separated on Euclidean distance be-tween log-scaled resource affinities and niche overlap (Fig. 4D). Given the technical challenges of quantifying resource affinities for cross-fed metabolites *in vitro*, we utilized our consumer-resource framework to identify more accessible signatures of robust richness. We found that species in the most rich (top 10%) communities were consistently able to grow well on the spent medium of their partners, in contrast to the least rich (bottom 10%) communities (Fig. 4E). We then conducted similar experiments *in vitro*: we cultivated each of the four species in monoculture to stationary phase on two different carbon sources, malate or mannose (for Ct we used isoleucine instead), to create spent media. These eight spent media were then used to cultivate the four species in monoculture, with growth compared against no-carbon controls. In agreement with the most diverse (top 10%) simulated communities, all four species exhibited robust growth on the metabolic byproducts of their partners (Fig. 4F). While self-facilitation was observed, as reported previously, ^18,34,54^ growth was more pronounced when species were cultivated on the spent media of other community members (Fig. 4F). Across all conditions, distributions of fold-change growth qualitatively mirrored the results from the top 10% simulated communities (Fig. 4E and 4G), providing empirical evidence that a network of cross-feeding interactions underpins the observed coexistence.

### Interaction patterns support extensive cross-feeding and metabolic niche sepa-ration

The sign and magnitude of pairwise species interactions are key determinants of the emergent properties of microbial communities. ^55–57^ To elucidate the association between interactions and coexistence, we quantified pairwise interaction scores by calculating the log-response ratio, defined as the log_10_ ratio of a species’ final abundance in co-culture relative to its abundance in mono-culture. ^57^ Under this framework, positive interaction scores indicate facilitation, while negative scores represent competition. In fresh, unsupplemented media, our community displayed a striking dominance of positive interactions (Fig. 5A, Table S1). Not surprisingly, in mannose (where only At can grow independently, Ct cannot utilize mannose and Oa and Ml are auxotrophs), all interactions were positive with the exception of those where At served as the focal species, as it provided the primary metabolic support for the community without receiving equivalent benefits in return. Supplementing the media with amino acids and/or vitamins weakened and shifted the interactions towards negative (Fig. 5A), mirroring findings in toxic industrial environments where nutrient supplementation similarly reduced positive interactions,^22^ suggesting that more benign environments reduce facilitation between species.

**Figure 5:**
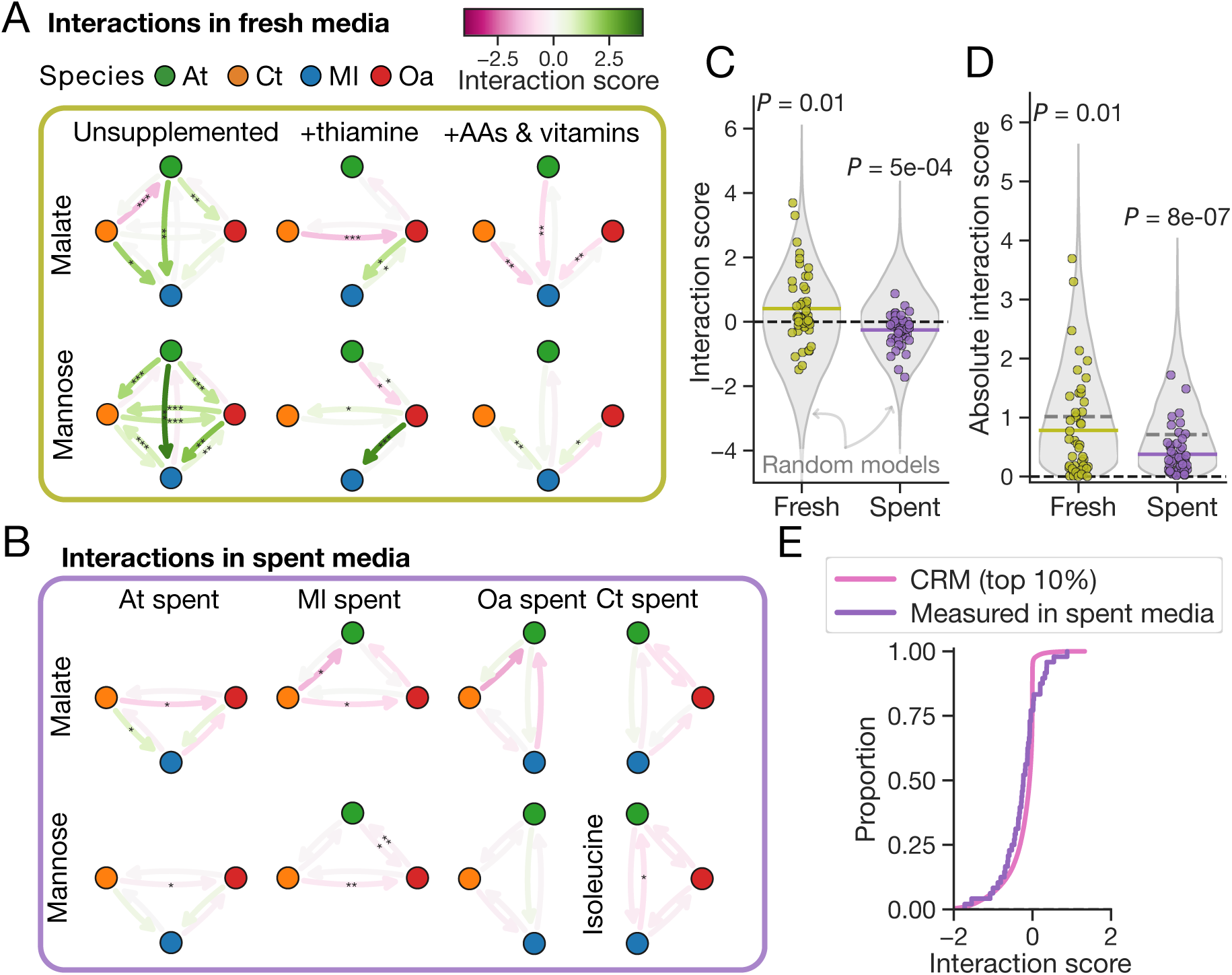
Species interactions in fresh and spent medium. A) Pairwise interaction scores in fresh malate and mannose (isoleucine) media, illustrating the shift in interaction strength and sign following vitamin and amino acid supplementation. B) Interaction scores in spent media with either malate, mannose or isoleucine as the original carbon source. Interactions were quantified as the log_10_ fold-change in population size (CFUs/mL) of co-cultures relative to mono-cultures. Statistical significance in A and B was determined on log_10_-transformed population sizes using Welch’s t-test with Benjamini-Hochberg correction (\**P* < 0.05, \*\**P* < 0.01, \*\*\**P* < 0.001). C) Interaction scores across fresh and spent media compared against null distributions from random models (Methods). Coloured horizontal bars indicate mean values. A two-sided one-sample Welch’s *t*-test was used to assess the significance of the mean deviation from zero. D) Magnitude of interaction scores compared against null distributions from random models (as in C). Colored and gray horizontal bars represent the median absolute scores of the experimental data and the null models, respectively. A two-sided one-sample Wilcoxon signed-rank test was used to assess whether the observed median magnitude significantly deviated from the null expectation. E) Cumulative distribution functions comparing *in vitro* and *in silico* (top 10%) interaction scores in spent media.

While interactions in fresh media elucidate dependencies and competition for the initial re-sources, they don’t resolve how species partition or compete over metabolic byproducts in the assembled communities. We reasoned that if the species effectively partition the available byproducts, they should exhibit weak interactions in spent media. Furthermore, if the vitamins and amino acids required by the auxotrophs were already abundant in the spent media, we expected fewer strong positive interactions. Indeed, we observed a significant shift towards negative interactions in spent media (Fig. 5B and C, Table S1). Interaction scores are fundamentally constrained by the yield of the growth environment, complicating the comparison of interaction strengths. Therefore, to estimate distributions of interaction scores expected by chance, we calculated interaction scores from 10,000 random mono- and co-culture population sizes from lognormal distributions fitted to the range of observed values in fresh and spent media, respectively (Methods). In comparison to these null distributions, absolute interaction strengths were lower than expected in both fresh and spent media, but the difference was pronounced in spent media (ratio of expected and observed median: 0.78 in fresh vs 0.54 in spent, Fig. 5D). Furthermore, the distribution of interactions in spent media aligned well with the distribution of simulated interactions in the top 10% rich communities (Fig. 5E). The consistent convergence of our experimental results with predictions from the most rich (top 10%) CRM communities suggests that our four-species community is an example of a community where metabolic interactions ensure robust coexistence across divergent environments.

## Discussion

Our results demonstrate that a synthetic four-species bacterial community can maintain robust, stable coexistence across a vast array of nutrient environments, regardless of external resource complexity. We rule out several alternative explanations and show that this robust coexistence can be explained by niche partitioning of metabolic byproducts from the co-cultivated species. It is already known that microbes create chemically rich exometabolomes,^46–48,58,59^ and that niche partitioning of such exometabolomes occurs in natural microbiomes. ^60^ Here we show that such niche partitioning is sufficient to support coexistence. Although the persistence of four species on a single carbon source may superficially appear to violate the competitive exclusion principle, this view is incomplete. Instead, our data suggest that coexistence emerges from intense niche creation, increasing the number of available resources and thus the theoretical limit on community richness. In light of previous work also showing coexistence of multiple species on a few provided resources, ^17,18,29^ we hypothesize that this phenomenon is the norm rather than an exceptional feature of our specific four-species community. It is important to note, however, that the extrapolation of these findings to natural communities requires careful consideration. As natural ecosystems typically contain hundreds or thousands of different species,^61,62^ it remains unknown whether the diversity of metabolic byproducts is sufficient to provide a stabilizing niche for every member of such highly complex assemblages.

A second central finding of this study is that robust coexistence appears to be an intrinsic feature of the community itself – ingrained in the species’ metabolic byproducts and the differences in their catabolic capacities. While metabolic versatility is a classic determinant of coexistence,^63,64^ our simulations suggest that variance in resource affinities plays a key role in maintaining richness across environments. In accordance with these simulations, experimental data show substantial variation in resource affinities within and between species^53^ and that such variation leads to niche partitioning of provided resources. ^65^ High-throughput quantification of resource affinities across diverse nutrients and species could further illuminate how natural variability in these traits enables coexistence. Such efforts may eventually allow for *in vitro* experiments with communities that span a broad range of affinity profiles. Our experiments indicate that our specific 4-species community has the hallmarks of consistently rich communities as predicted from CRMs. Some of these hallmarks, such as affinities for cross-fed metabolites, might be cumbersome to measure and might not scale to the design of more complex synthetic communities. In contrast, spent medium experiments are straightforward and potentially equally informative. As the design of synthetic communities becomes increasingly relevant for human health, agricultural or industrial applications, these findings suggest a feasible avenue towards designing communities whose coexistence remains robust to the variation in such environments.

## Acknowledgments

We thank all members of the Mitri lab at the University of Lausanne for discussions and feedback.

## Author contributions

Conceptualization: SS, SM; Methodology: SS, MT, EU, PP, ST, PP, DM, SM; Investigation: SS, MT, EU, AG, PP; Visualization: SS; Funding acquisition: SS, SM; Project administration: SS, SM; Supervision: SS, SM, DM; Writing – original draft: SS; Writing – review & editing: All authors.

## Data and materials availability

The strains used in this study are available on request. All data and code used for simulations and data analyses are available on GitHub at https://github.com/Mitri-lab/coexistence. A permanent archive of this repository will be deposited to Zenodo upon publication.

## Methods

### Strains, media and culturing conditions

All experiments were conducted with one or more of these four strains previously isolated from and cultivated in metal-working fluids: *Agrobacterium tumefaciens* MWF001 (At), *Comamonas testosteroni* MWF001 (Ct), *Microbacterium liquefaciens* MWF001 (Ml; now classified as *M. maritypicum*) and *Ochrobactrum anthropi* MWF001 (Oa; now classified as *Brucella anthropi*). ^22,31^ These species can be distinguished on selective media and quantified by plating and counting of colony-forming units (CFUs).^32^

Most experiments were conducted in M9 minimal medium with different carbon sources. Unless otherwise stated, the amount of carbon in each medium was normalized to 90 mM of carbon atoms. The different M9 minimal media were prepared by mixing 10 mL of autoclaved 10X M9 salts (Sigma-Aldrich, US, M6030-1KG), 1 ml of 50X HMB, ^35^ 5 mL of 10X carbon source stock and 34 mL of MilliQ water. Finally, all media were pH-adjusted to pH 7.4 (with NaOH / HCl) and sterile filtered.

For all experiments, strains were streaked from glycerol stocks onto Tryptic Soy Agar (TSA) and incubated at 28°C for 2 days. Single colonies were used to inoculate 10 mL Tryptic Soy Broth (TSB) in 50 mL Erlenmeyer flasks and grown overnight. These precultures were then diluted to an OD_600_ of 0.05 in 20 mL fresh TSB in 100 mL flasks and incubated for 3 h (28°C, 200 rpm). Prior to inoculation of experimental assays, 10 mL aliquots of these cultures were washed by centrifugation (3220 rcf, 6 min), the supernatant removed, and the pellet resuspended in 1 mL carbon-free M9 minimal medium. The resuspended cells were centrifuged again (8000 rcf, 4 min), the supernatant discarded, and the pellet resuspended in 1 mL carbon-free medium. This last wash step was repeated once.

### Growth phenotyping

We characterized the metabolic repertoire and dependencies of the four species through cultivations in M9 minimal media. Previous laboratory studies identified Oa as a thiamine auxotroph on solid media. Additionally, the inability to cultivate Ml in monoculture suggested that this species was also an auxotroph.

We first verified that Ml was cultivable in 15 mM glucose M9 medium when supplemented with a vitamin mix composed of biotin (Acros organics, D(+)-Biotin, 96%), thiamine hydrochloride (Fluka, Vitamin B1 hydrochloride #95160/25g; herein referred to as thiamine) and niacin (Sigma-Aldrich, Nicotinic acid #72309/100g) and an amino acid mix composed of all 20 proteinogenic amino acids (Table S2). Vitamins were added to final concentrations of 0.1 mg/L biotin, 0.5 mg/L thiamine and 1 mg/L niacin. All amino acids were supplemented to a final concentration of 0.15 g/L each. We initially used GapMind^66^ to identify potential amino acid auxotrophies in Ml. GapMind flagged histidine, lysine, tyrosine and tryptophan as potential auxotrophies for Ml, but supplementing the glucose M9 medium with the vitamin mix and these four amino acids did support growth of Ml. We therefore conducted an amino acid dropout screen in glucose M9 plus vitamin mix medium featuring all 20 N-1 proteinogenic amino acid combinations in addition to all amino acids and no amino acids controls. Following the previously described preculturing protocol, Ml was inoculated to OD_600_=0.01 in 4 mL culture medium in straight glass tubes (15.8 cm tall, 1.5 cm diameter) for 114 hours (28°C, 200 rpm, two replicates). Growth was monitored by measuring OD_600_ directly in the glass tubes at nine time points spread over the course of the experiment (NANOCOLOR VIS II, Macherey-Nagel, Düren, Germany). The N-1 amino acid dropout experiment revealed proline and cysteine as potential auxotrophies (Fig. S1D). To verify these auxotrophies, we conducted a follow-up experiment with supplementation of proline, cysteine or both (Fig. S1E, three replicates, 188 hours, otherwise similar experimental procedure, equipment and parameters). In the same experiment, we also verified the vitamin auxotrophies of Ml and Oa that were predicted from the presence/absence of key genes (according to ^67^) in the annotated genomes obtained from NCBI under the accession code PRJNA991498^23^ (Fig. S1A). In agreement with the absent genes, the growth of Ml was reduced when leaving out either biotin or thiamine, and the growth of Oa increased substantially on thiamine supplementation.

To select a set of carbon sources for further growth phenotyping, we used a computational approach to identify carbon sources that were metabolically diverse. For this purpose, we used the universal genome-scale metabolic model (GEM) for bacteria from CarveMe. ^68^ Using this GEM, we predicted intracellular flux distributions with 509 different metabolites using parsimonious FBA^69^ with growth as the objective. This optimization was conducted using COBRApy^70^ and Gurobi (v10.0.3), setting bounds on exchange reactions to mimic aerobic growth in a minimal M9-like medium (Table S3). The uptake rate of the carbon source was set to 10 mmolgDW^-1^h^-1^. To identify groups of metabolites with different flux patterns we first converted fluxes to binary values (0, -1 or 1), discarded fluxes with zero variability, conducted a Principal Component Analysis (PCA) on the remaining binary flux matrix, and finally used HDBSCAN^71^ to group metabolites based on principal components 1 and 2 (Fig. S2A). We then selected 33 carbon sources that differed in metabolite class (amino acids, organic acids, simple sugars, alcohols, nucleoside and others) and covered all 9 clusters (Table S4). We then tested the ability of the 4 species to grow on each of these carbon sources in minimal M9 medium, using concentrations corresponding to 90 mM carbon atoms (except for Ct in citrate, where we used 5 mM rather than 15 mM citrate because of Ct’s long lag-time in high citrate concentrations). Additionally, for Oa we supplemented the media with 0.5 mg/L thiamine, and for Ml we supplemented the media with 0.1 mg/L biotin, 0.5 mg/L thiamine, 0.18 g/L cysteine and 0.1 g/L proline. We also included a no-carbon-source control condition (which also included the vitamin and amino acid supplements). For these experiments, each strain was precultured and washed as described above and then inoculated to OD_600_=0.05 in 200 μL of media in 96-well plates and cultivated with continuous shaking (double orbital, 425 cpm) in a BioTek Synergy H1 (Agilent Technologies, Winooski, VT, USA) plate reader at 28°C for 72 hours (Fig. S2B).

We quantified the ability of each species to grow on each of the 33 carbon sources qualitatively (growth or no growth) and quantitatively (maximum growth rate, lag-time, yield). A carbon source was classified as growth-supporting if cultures exceeded a minimum yield (OD_max_ - OD_start_ > 0.05) and if this yield was statistically greater than the no-carbon control (*P* < 0.05, Welch’s one-sided *t*-test, N = 3). Growth parameters were extracted from the growth curves using curveball, ^72^ but only fitting growth models until the time of maximum OD_600_. Regardless of which model curveball selects as the best fit, the common parameters are: maximal growth rate (*µ_max_*), the lag time (*t_lag_*), the final yield (*Y*) and the initial starting OD (*y*_0_). From these four parameters, we define the effective growth rate (*µ_eff_*) as the constant exponential rate that would be required to reach the observed yield *Y* from the initial density *y*_0_ in the total time. In the absence of lag, the effective growth rate and the maximal growth rates are equal (at high resource affinity levels). With nonzero lag time, the effective growth rate is given by

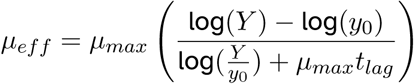

This measure thus captures the joint effect of lag time and maximal growth rate on competitive ability, rather than relying on *µ_max_* alone. This is important since the relationship between these parameters has been shown to impact competitive ability. ^41^

### Community assembly experiments

The community assembly experiment was conducted in 32 different M9 minimal media, with 16 single carbon sources, 15 pairs of carbon sources and a no-carbon control environment (Fig. 1). The four species were precultured and washed as described above. Washed cultures were combined at equal densities (each species to OD_600_ = 2), vortexed and used to inoculate all wells of a 96-well plate to an initial OD_600_ = 0.03 (in 200 μL) of each species to start the first growth cycle. Each transfer of the community assembly was cultivated for 72h in a plate reader (Biotek, Synergy H1) at 28°C with continuous shaking (double orbital, 425 cpm). On each transfer a 1/100 dilution (2 μL) was performed into a freshly made plate. However, wells with citrate or inosine as the carbon source were only diluted 1/10 because of low yield (first transfer of citrate wells were 1/100). All wells were carefully mixed using a 200 μL multiwell pipette set to 70 μL before each transfer. The abundance of each species at the end of each growth cycle was quantified by counting CFUs on selective media. ^32^ Quantified population sizes (as CFUs/mL) were log_10_-transformed prior to statistical analyses and calculation of abundance ranks. Log_10_-transformed CFUs from the two last transfers were used to compute final abundance ranks (by comparing means) and for statistical analyses. To compare the final abundance of each species in each condition with the final abundance in the no-carbon control we used the two-sided Welch’s *t*-test for At, Ct, and Oa, and the one-sample one-sided *t*-test for Ml with respect to the limit of detection. The limit of detection was set to one CFU in the undiluted 5 *µ*L volume used for CFU quantification, i.e. 200 CFUs/mL. For the second community assembly experiment (Fig. 2), six single carbon source environments (acetate, glutarate, L-glutamate, L-histidine, malate, ribose) were repeated exactly as in the first community assembly experiments. We also included 18 conditions with the same carbon sources but either at different carbon source concentrations (5X: 450 mM and 1/50X: 1.8 mM carbon) or supplemented with vitamins (0.1 mg/L biotin, 0.5 mg/L thiamine) and the amino acids required by Ml for rapid growth (0.15 g/L proline, 0.15 g/L cysteine). All conditions were conducted in triplicates. For the second community assembly experiment, the quantified population density after the sixth transfer was used in three cases as the final population size because of issues with the CFUs from transfer 7 and 8 (two wells for Oa in Inosine 1/50x, one well for Ml in LB).

### Community assembly experiment in chemostats

Chemostat experiments were performed using Chi.Bio reactor modules^73^ coupled to an Ismatec IPC multichannel peristaltic pump. Cultures were grown in 30 mL glass vials (Fisher Scientific, 11593532) sealed with 20 mm septa secured by hole caps (Supelco, 22 mm). Four ports were punched into each septum and fitted with PTFE tubing (ID 1/16 in, OD 1/8 in; Altec 01-96-1702) to provide lines for medium inflow, effluent outflow, sampling, and aeration/pressure release. Fresh medium was supplied from a 0.5 L glass reservoir to the growth vial using 2-stop Tygon E-Lab tubing (ID 0.76 mm) and silicone pump tubing (bore 2.5 mm, wall 1 mm; Altec 01-93-1416/20). Effluent was removed via a PTFE outflow line connected to silicone pump tubing (bore 2.5 mm, wall 1 mm) placed directly in the peristaltic pump and routed to a waste bottle. The working volume was maintained at 20 mL by fixing the height of the PTFE outflow tube to act as an overflow at the 20 mL fill level. Cultures were aerated using an aquarium air pump (JBL ProAir a50) connected to the aeration port through an inline air filter and mixed continuously with a magnetic stir bar (10 mm; Fisher Scientific 11732513). After assembly (vial, tubing, and feed/waste bottles), the complete setup was sterilized by autoclaving prior to inoculation. The peristaltic pump was programmed to a flow rate of 33 *µ*L min*^−^*^1^, corresponding to a dilution rate of *D* = 0.1 h*^−^*^1^ for a 20 mL working volume, and reactors were maintained at 28 *^◦^*C.

Five minimal-medium conditions were prepared, each standardized to a total of 90 mM carbon atoms, using ribose, sodium acetate, histidine, or glutaric acid as the sole carbon source, as well as a no-carbon control. At, Ct, Ml, and Oa were grown as described above. Prior to inoculation, cultures were adjusted and combined to assemble four-species communities with equal starting abundances and a total initial density of OD_600_=0.03 in each medium condition. Chemostats were inoculated on day 0 with the prepared community. From day 1 to day 4 after inoculation, cultures were sampled twice daily by withdrawing 700 *µ*L through the septum, preparing 10-fold serial dilutions, and spot-plating 5 *µ*L drops of each dilution on selective agar for counting CFUs of each species (see above).

### Mono- and co-cultures in fresh and spent media

Spent media were created in the following way: The four species were precultured and washed as described above. Then, the species were inoculated (in monoculture) to OD_600_=0.05 in 10 mL of the M9 media with glutamate, mannose, glycerol, isoleucine, or malate and cultured for 72h (50 ml Erlenmeyer flask, 200 rpm, 28°C). Additional thiamine was included in Oa media, and additional thiamine, biotin, proline and cysteine were included in Ml media (concentrations as in the second community assembly experiment). Optical densities were measured at 70h and 72h to ensure that the cultures had reached stationary phase (OD_600_ readings were unchanged between these time points). The spent culture media were filtered (0.22 µm pore size) and stored in falcon tubes at -20°C until use. Sterility was verified by plating filtered media on TSA (28°C, >48h); any contaminated batches were refiltered and re-tested prior to use.

Mono- and co-cultures in fresh and spent media were conducted in the following way: The four species were precultured and washed as described above, and then inoculated (either in monoculture or in pairs) to OD_600_=0.001 (for each species) in 200 µL of spent media in 96-well plates (see Table S5 for an overview of all combinations). All experiments were conducted in triplicates. The experiment with cultivations in spent media based on mannose/isoleucine was repeated once because of issues with certain CFUs, and our data include data from both experiments. Inoculum sizes were also quantified as CFUs. The cultures were incubated in a plate reader (Biotek, Synergy H1) at 28°C for 72 hours with continuous shaking (double orbital, 425 cpm). Final populations of the different species were quantified as CFUs by serial dilutions and plating on selective media as described above.

### Quantification of interactions

Interactions in fresh and spent media were quantified as the log-response ratio ^57^ and the significance of each interaction was assessed using the two-sided Welch’s *t*-test to compare CFUs in mono and coculture. CFUs were log_10_ transformed prior to quantification of interaction strengths and significance. Thus, the interaction (*Y_i_j*) measuring the effect of species *j* on focal species *i* was calculated for each condition and for each partner *j* as 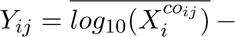 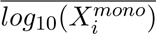, with 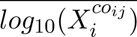 and 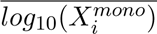 denoting the mean of log-transformed CFUs in co-culture and mono-culture, respectively.

To quantify expected distributions we used random models to create null-distributions of interaction scores in two ways: 1) We fitted normal distributions to match the 5% and 95% percentiles of the log_10_-transformed measured CFUs in fresh and spent media, respectively. From each distribution, we sampled 10,000 pairs of random log_10_-transformed CFU values (representing final abundance in mono- and co-culture) and calculated the corresponding distribution of interaction scores. 2) We sampled with replacement 10,000 pairs of log_10_-transformed CFU values from the pool of measured values (for fresh and spent media separately), and calculated the distribution of interaction scores from these pairs. For the statistical comparisons of the magnitude of interactions scores, both approaches gave similar results and the same conclusion. The measured and fitted distributions of log_10_-transformed CFUs and a comparison vs measured interactions and null distributions from the second approach are shown in Fig. S6 (corresponds to Fig. 5C and D).

### Consumer resource model

The consumer resource model presented in this paper is based on previous modifications of Robert MacArthur’s consumer resource model^50,74^ that extends this framework to include cross-feeding.^18,29,52^ Briefly, the framework models the abundance of both species and resources, and the growth of each species is a function of its nutrient preferences, resource concentrations and metabolite release.

We denote the set of species as *N_i_* where *i* = 1*, …, N*, and the set of resources as *R_j_* where *j* = 1*, …, M*. Each species is characterized by a resource uptake matrix *J_ij_* which represents the rate at which species *i* consumes resource *j*, leakage parameter *l* that specifies how much of the consumed resources are released back into the environment, a metabolite release matrix *D_ijk_* that describes the transformation of resource *j* to resource *k* for species *i* upon metabolite release, a yield parameter *g_i_* and a maintenance parameter *m_i_*. The entries in *J_ij_* are calculated from a maximum uptake rate matrix *C_ij_*and a substrate affinity matrix *K_ij_*according to the Monod equation: ^75^

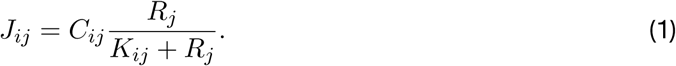

Each resource is described by an energy content *w_j_*. The per capita growth of each species is then described by:

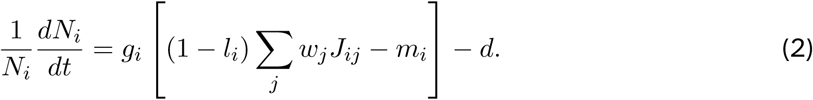

Here, *d* is the dilution rate of the system. Resource concentrations are then described by the following equation:

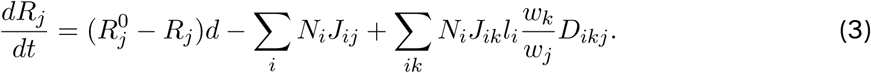

Here, 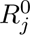 represents the initial carbon source concentration of resource *j*. For all simulations we assumed that resources had the same energy content (*w_j_* = 1), neglected energy used for maintenance (*m_i_*= 0) and assumed an equal yield (*g_i_*) of 0.1 for all four species.

Entries for the matrices *C*, *D* and *K* were sampled from random distributions using scipy. ^76^ For *C*, we sampled maximum uptake rates from a lognormal distribution fitted to the distribution of maximum growth rates (*v_max_*), scaled to uptake rates (*c_max_*) based on a yield (*g*) and leakage fraction (*l*) of 0.1 (*c_max_* = *v_max_*/*g*(1 − *l*)) on single carbon sources in monoculture (Fig. S2B). Hence, *C_ij_* ∼ LogNormal(*µ, σ*), where *µ* = 0.37 and *σ* = 0.91 are the mean and standard deviation of the log-scaled data, respectively. To reduce the number of simulations where all species were diluted out, we defined a minimal threshold for the maximum uptake of the single, initial carbon source (*j* = 0) across the four species, i.e.

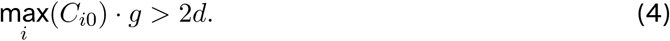

To create *D*, we first sampled values from a Uniform distribution (*D_ijk_* ∼ Uniform(0, 1)). Then, we set the diagonal elements to zero to avoid a resource being leaked as the same resource (*D_ijj_* = 0, ∀*i* ∈ *N,* ∀*j* ∈ *M*). Finally, we normalized the rows of each species’ *D* matrix to ensure energy conservation (Σ_k_*D_ijk_*= 1, ∀*i* ∈ *N,* ∀*j* ∈ *M*).

### Simulating four-species microbial communities

We simulated continuous cultivations of four random species for 2000 time steps (a.u.) at a dilution rate of 0.1, with an initial abundance of each species equal 10*^−^*^3^ (a.u) and a single resource with an initial concentration 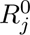 of 10 (a.u.). We determined the convergence of the simulation based on the change in species abundance in the last timestep. If no species abundance changed more than 10*^−^*^7^ in the last time step we considered the simulation as converged. Simulations that did not converge, or failed due to numerical issues, were discarded.

We simulated four-species communities with different levels of niche breadth and different levels of variation in resource affinities. We define niche breadth as the fraction of non-zero entries in *C_ij_* for each species *i*. Non-zero entries in *C_ij_* were selected randomly according to the given niche breadth (*P* (*C_ij_* = 1) = niche breadth). Resource affinities *K_ij_* were sampled from a lognormal distribution (*K_ij_* ∼ LogNormal(*µ, σ*)), with fixed mean *µ* = −1 (in log-space) and varying *σ*.

