## Supplemental Figures for "Cross-feeding enables robust coexistence between four bacterial species"

### Supplementary Figures

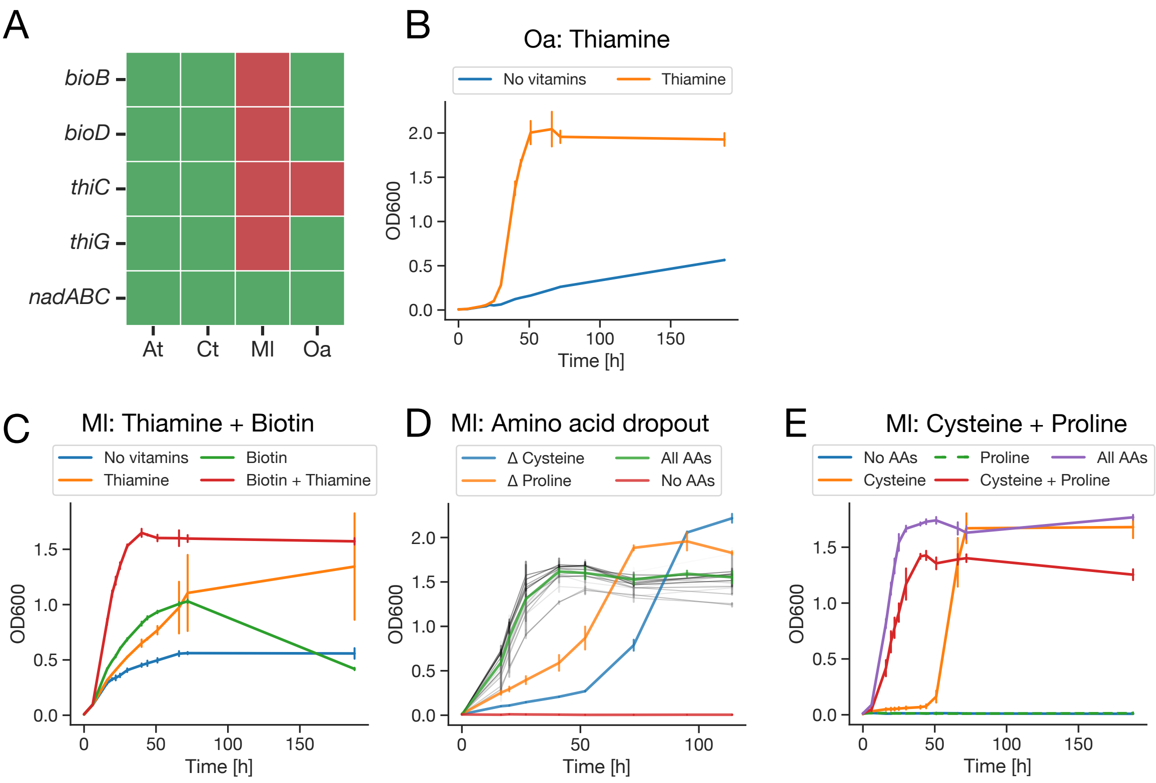


Figure S1: A) Presence/absence of key genes for biotin (*bioB* and *bioD*), thiamine (*thiC* and *thiG*) and niacin (*nadA*, *nadB* and *nadC*) biosynthesis in the four species based on the annotated proteome. B & C) Evaluating vitamin auxotrophy predictions based on proteome annotations of Oa and Ml. Thiamine was needed to support rapid growth and high yield of Oa. Both biotin and thiamine were needed to support full growth rate and yield for Ml. All media here was based on a glucose M9 medium, and for the Ml cultivation all 20 proteinogenic amino acids were supplemented. D) Amino acid dropout screen to identify the amino acid auxotrophies of Ml. Ml only showed reduced growth in the Δproline and Δcysteine conditions. Other dropouts are shown in light gray. All media here, and in panel C, was based on a glucose M9 medium supplemented with biotin, thiamine and niacin. E) Confirmation of Ml auxotrophies identified in amino acid dropout screen. While external supply of cysteine is strictly needed for Ml's growth, the lack of proline induces a 2-day lag-time followed by rapid growth.

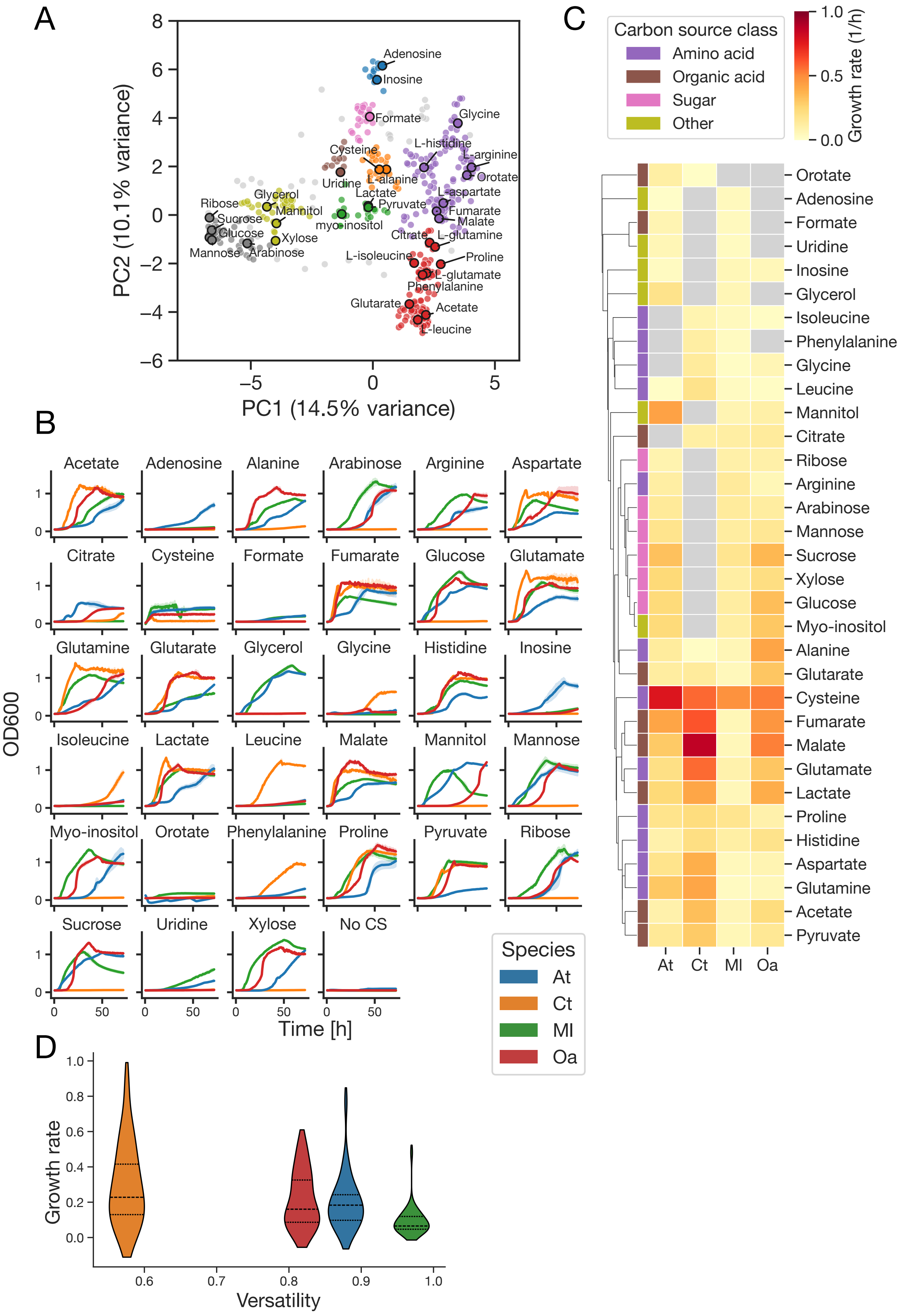


Figure S2: Growth phenotyping on 33 carbon sources. A) A PCA used to identify metabolically diverse based on differences in predicted flux patterns (see Methods). By selecting carbon sources from all of the nine clusters we covered metabolic variability. Annotated carbon sources are those selected to be used for growth phenotyping. B) Growth curves of each species in monoculture for each of the 33 carbon sources. Monocultures of Ml and Oa were supplemented with essential vitamins and amino acids (see Methods). C) Estimated maximum growth rate

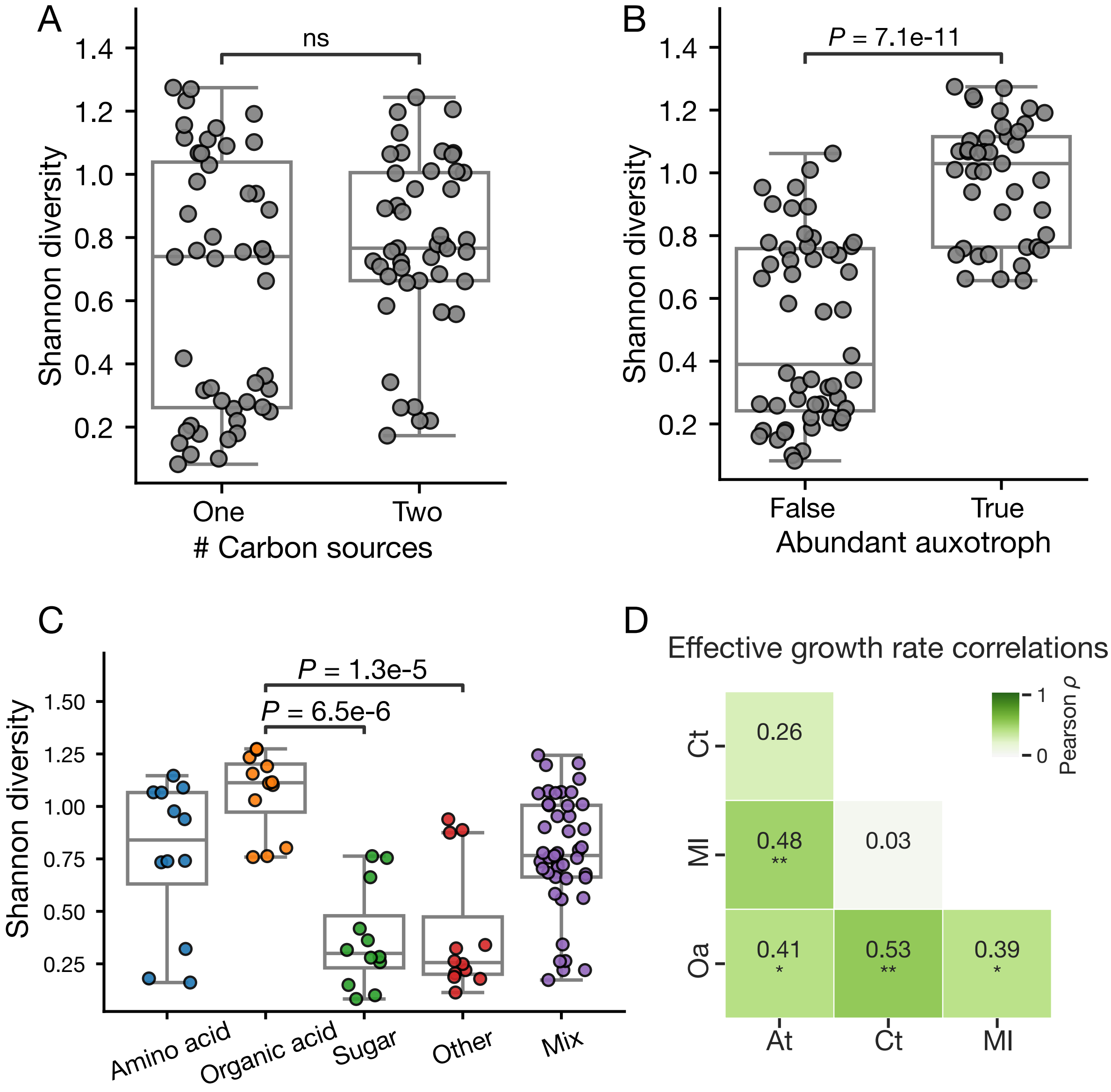


Figure S3: Shannon diversity of community assembly experiment and growth rate correlations. A) We find no significant difference in Shannon diversity when we simply group by the number of carbon sources ($P = 0.21$, Mann-Whitney U). B) We find however that communities where one of the auxotrophs comprise > 10% of the total abundance to have a significantly higher Shannon diversity ($P$ = 7.1e-11, Mann-Whitney U). C) We find also significant differences in Shannon diversity between types of carbon sources. Significance assessed using the Kruskal-Wallis test followed by Dunn's posthoc test with Benjamini-Hochberg correction. D) Pearson correlation between effective growth rates (*$P < 0.05$, **$P < 0.01$).

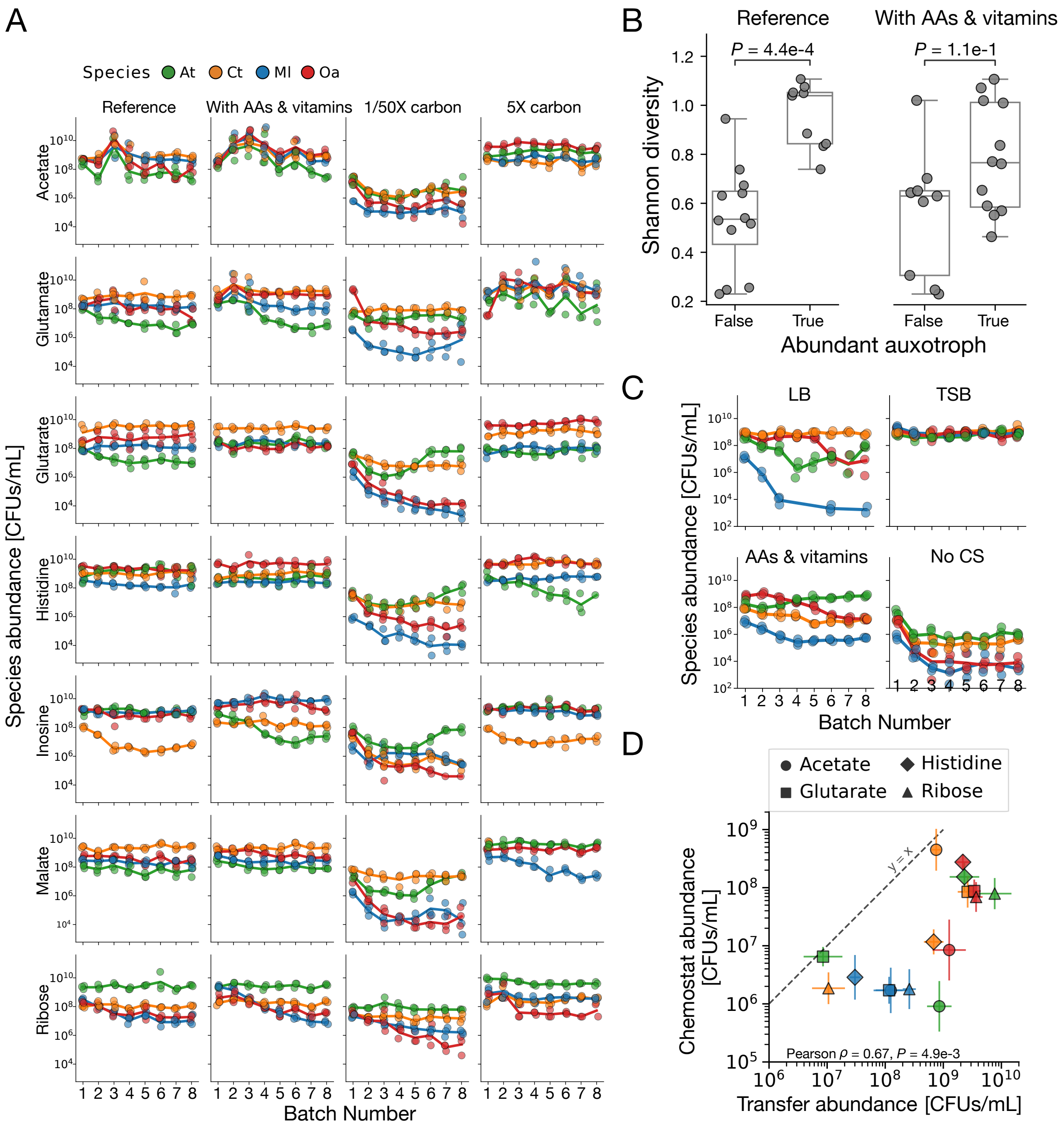


Figure S4: Community-assembly experiments to test alternative hypotheses. A) Quantified species abundance at the end of each of the 8 consecutive batch transfers in the reference condition, with vitamin and amino acid supplementation, with reduced and increased amount of carbon. B) A comparison of community assembly endpoint Shannon diversity with or without an abundant auxotroph (Ml or Oa >25% of the population) in the reference conditions and when supplemented with vitamins and amino acids. C) Quantified species abundance at the end of each of the 8 consecutive batch transfers in LB, TSB, in the amino acid and vitamin supplement, and in the no-carbon control. D) A comparison of final population sizes in chemostats and in the transfer experiments. The mean and standard deviation is calculated from the (2/3) samples after 41 hours for the chemostats and from the last two transfers for the transfer experiment. The Pearson correlation is calculated from the $\log_{10}$-transformed mean values.

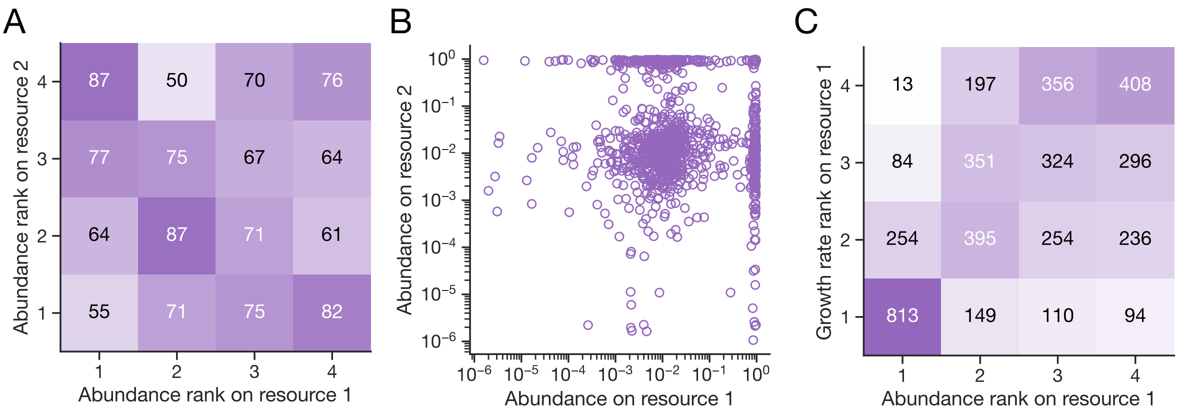


Figure S5: Comparisons of abundance, abundance ranks and growth rates on resource 1 and 2 for consumer-resource models. A) No correlation between the abundance rank on resource 1 and 2 (Kendall's $\tau$ = -0.05, *P* = 0.06, N = 1132). B) A very weak, negative correlation between $\log_{10}$-transformed abundances on resource 1 and 2 (Pearson $\rho$ = -0.06, $P$ = 0.04, N = 1132). C) In accordance with our experimental results, the growth rate rank and abundance rank is positively correlated in our consumer-resource model simulations with four coexisting species (Kendall's $\tau$ = 0.45, *P* = 1e-304, N = 4664)

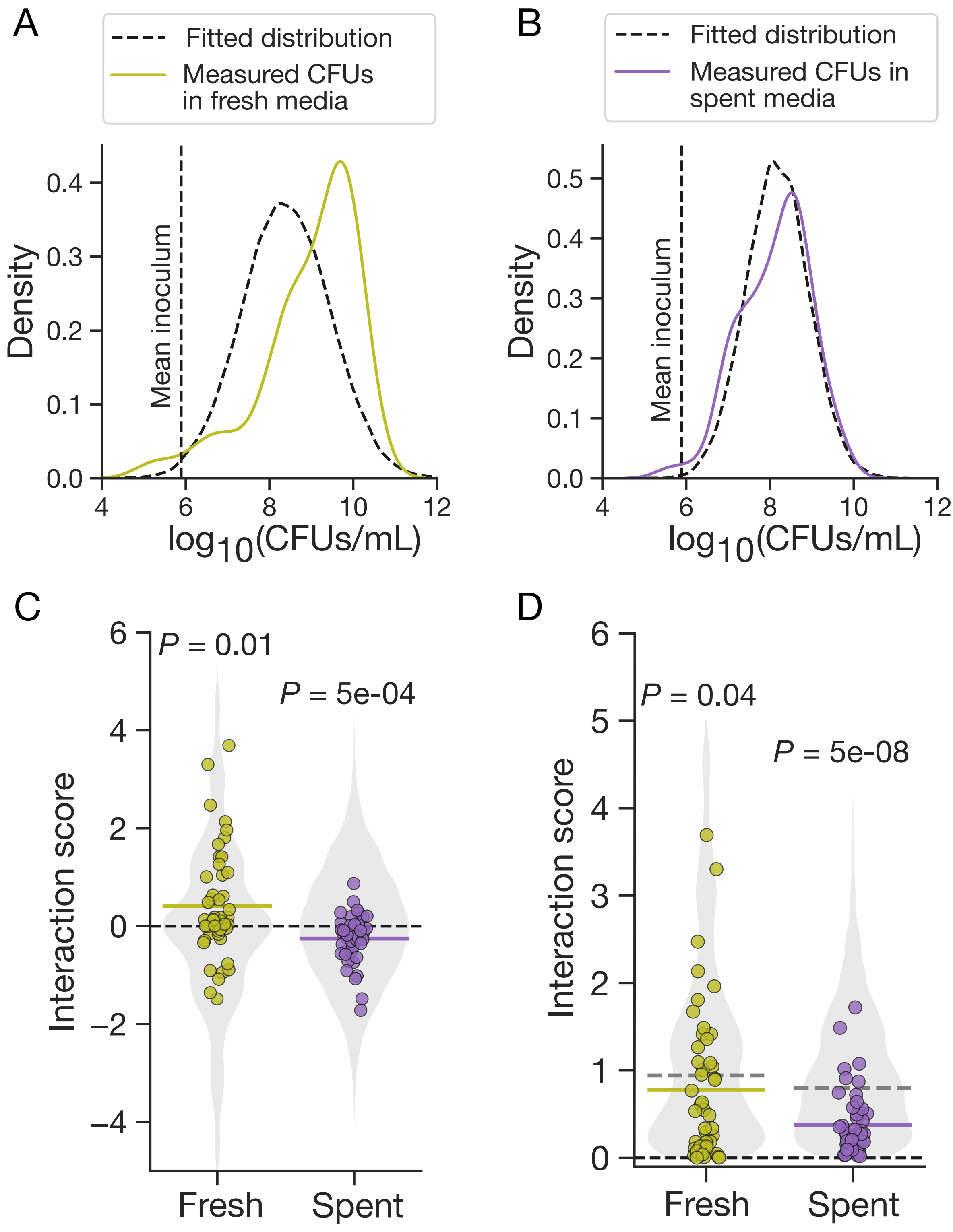


Figure S6: Random models of interaction scores. A and B) Distributions of $\log_{10}$-transformed CFU values and lognormal distributions fitted to 5% and 95% percentiles of measured interactions, used for comparisons if Fig. 5. C) Interaction scores across fresh and spent media compared against null distributions from random models based on permutations of measured CFU values (Methods). Coloured horizontal bars indicate mean values. A two-sided one-sample Welch's $t$-test was used to assess the significance of the mean deviation from zero. D) Magnitude of interaction scores compared against null distributions from random models (as in C). Colored and gray horizontal bars represent the median absolute scores of the experimental data and the null models, respectively. A two-sided one-sample Wilcoxon signed-rank test was used to assess whether the observed median magnitude significantly deviated from the null expectation.
